# Nanopore-based DNA sequencing in clinical microbiology: preliminary assessment of basic requirements

**DOI:** 10.1101/382580

**Authors:** Håvard Harstad, Rafi Ahmad, Anders Bredberg

## Abstract

**Aim:** Identify basic requirements for a metagenomic nanopore sequencing protocol permitting frequent application in a clinical microbiology daily routine diagnostic setting.

**Background:** Nanopore sequencing with the Oxford Nanopore Technologies MinION device has a potential to markedly improve clinical diagnosis of infections. Reports have emerged recently that it may provide direct-from-clinical-sample information; for example, with urine samples, bronchial tuberculosis samples and orthopedic prostheses. However, the ideal protocol for clinical use remains to be determined, especially relating to detection of relevant pathogen quantities and to finding a reasonable level of economic costs.

**Results:** MinION can provide qualitatively and quantitatively correct identification of multiple species in metagenomics samples. For detection of clinically relevant quantities of bacteria (on a nanogram DNA level) there is a need for carrier DNA. Importantly, high-purity DNA and a naïve MinION flow cell seem to be critical parameters.

**Conclusions:** Our results suggest that high-purity clinical sample DNA, addition of carrier DNA and a naïve flow cell are critical factors for clinical use of MinION. A relatively high error rate may limit detection of antimicrobial resistance genes, and a realistic level of costs will require availability of a price-reduced and single-use flowcell.

## Introduction

Nanopore-based metagenomic sequencing with the MinION device is a promising new tool for clinical diagnostic work (1-2). With the nanopore technique, you read the bases in a DNA strand as it threads through a nanometer-sized pore, by detecting changes in an ionic current across the pore as DNA bases pass through. Because it does not require DNA to be chopped into small pieces, as most other sequencing technologies do, you can get longer reads, which can be assembled more quickly into a full sequence. Another advantage over other metagenomics techniques is the real-time mode of analysis. For both RNA and DNA pathogens, the sequencer produces an immediate readout of the strand going through the pore (3).

This new technology will most probably become a standard part of routine clinical work for rapid identification and characterization of microbial isolates. The great potential of the technique has been described in Science ‥… it will make a tremendous difference both to diagnostics and to science it will quickly make its way into many labs and the clinic’ (4).

As illustrations of its clinical usefulness in threatening situations, a hemorrhagic fever virus (Ebola) was identified within 6 hours (5), and same-day diagnostic results for tuberculosis has been reported with the nanopore MinION device (6). The method has also been used for identification of viruses in a clinical sample (7), for bacterial pathogens in urine (8), in orthopedic prostheses-related tissue (9), in a liver abscess (10), in faeces (11) and to analyse for the anthrax agent (12) and to predict antibiotic resistance pattern in gonorrhea (13). Nevertheless, this clinical research field is quite new, and, as yet, the available studies have been on limited groups of pathogens and on few samples.

The aim of this pilot project was limited to exploring the possibility to use nanopore metagenomic sequencing for cost-beneficial, rapid and at informative diagnostic work in clinical microbiology.

## Methods

### Microbiology

The following reference bacterial strains were obtained from the American Type Culture Collection (ATCC): *Escherichia coli* ATCC 25922, *Klebsiella pneumoniae* ATCC 700603 expressing extended spectrum beta lactamase (ESBL) resistance (14), *Pseudomonas aeruginosa* ATCC 27853, *Proteus mirabilis* ATCC 29906 *and Haemophilus influenza* ATCC 49766. *Acinetobacter baumanii* is an in-house reference strain, Matrix Assisted Laser Desorption Ionization Time of Flight (MALDI-TOF MS microflex LT from Bruker Daltonics, Bremen, Germany) was used, with scores >2 verifying the strain identity before analysis.

### DNA extraction and sequencing

DNA was extracted with a robot employed by our clinical diagnostic laboratory for PCR analysis of patient samples (Roche MagnaPure Compact), or manually with standard silica mini-columns (Qiagen Genomic-tip 20/G) following the manufacturer’s instruction. DNA purity and concentration were determined using Nanodrop (NanoDrop 2000, ThermoFisher Scientific). Approximately 400-500 ng DNA was taken to build a DNA library for nanopore sequencing using a Rapid Sequencing Kit (SQK-RAD003 from Oxford Nanopore Technologies, Oxford, UK (ONT)) as described by the manufacturer, and then loaded onto a MinION sequencing apparatus flow cell. The MINION sequencing was performed with R9.4 flow cells (FLO-MIN106, ONT).

### Bioinformatics analysis of the obtained metagenomic sequences

The MinION sequencing was controlled using ONT’s MinKNOW software. The version of the software used varied from run to run but can be determined by inspection of fast5 files from the data set. Raw sequence reads were basecalled in real time via MinKNOW producing .fastq files. Basecalled data were then uploaded to the Epi2Me interface, a platform for cloud-based analysis of MinION data. Two different applications in Epi2Me were applied for data interpretation, What’s In My Pot? (WIMP) for identification of sequence data, and Antimicrobial Resistance Application (ARMA) software for identification of resistance genes.

In a number of instances, MinION sequencing control was shifted to customized MinKNOW scripts. These scripts provided enhanced pore utilization/data yields during sequencing, and operated by monitoring and adjusting flow cell bias-voltage (−180 mV to – 250 mV). More detailed information on these scripts can be found on the ONT community website. It is also worth noting that both software applications and chemistry in reagent kits and flow cells are under continuous development at ONT and updates appear frequently. As an example, the sequencing kit used in this report, SQK-RAD003, has been replaced by SQK-RAD004 having an updated composition. Furthermore, software updates often tends to imply changes in the interface + resetting all run scripts (if edited).

## Results and Discussion

Our first experiments followed a recently published clinical protocol for urinary tract infections (8). A urine sample submitted to our clinical laboratory and reported to the submitting clinician to have a clinically significant quantity (>10^8^/L) of *P. mirabilis*, as determined by our current standard diagnostic methods with Petri dish agar medium culturing and MALDI-TOF species identification, was used. After the sample had been anonymized, DNA was extracted on the Magnapure Compact in accordance with the published protocol. The purity and yield of the extracted DNA did not meet the requirements stipulated by the library SQK-RAD003 kit, but nevertheless a library was made. MinION sequencing showed dominance (77%) of eukaryotic (and presumably human) reads and 10% *P. mirabilis* reads. Furthermore the experiment produced very little data: 650 kb were produced within 2.5 hr. Because the results obtained by us with this protocol did not correspond to our needs for bacterial quantity sensitivity or DNA purity, and because of the low fraction of bacterial reads, we decided to focus on more basic protocol components involving DNA quality, MinION device and ONT software. All following experiments were performed without clinical samples, and instead with defined counts of bacterial reference strains, suspended in saline, and presumed problems associated with human samples such as DNA-degrading inflammatory substances and large quantities of patient DNA (15) were postponed until a later developmental phase.

Bacterial cultures at an 0.5 McFarland density, corresponding to approximately 10 bacterial cells/ml, of *E. coli* and *K. pneumoniae*, respectively, were mixed and DNA was extracted using both the MagnaPure Compact robot and manually with Qiagen Genomic-tip 20/G mini-columns. The results clearly show that the mini-column extraction is superior; there were more of informative reads and greater fragment lengths, obtained within a shorter real-time after loading onto the MinION flow cell (Table 1). For *E. coli* the fraction of informative reads was 48% and 92%, with Magnapure and mini columns, respectively; and the corresponding values for *K. pneumoniae* were 14% and 92%. Another observation was the longer read fragments with the mini-columns, likely to be due to gentle handling of the DNA during extraction.

**Table 1.** Sequencing results for a mixed *E. coli* and *K. pneumonia sample*, with two different DNA isolation protocols.

| Microorganism | DNA isolation method | MinION run time (hr) | Reads, informative (total) | Total yield (Mbp) | Best hit (reads) | Average length (kb) |
| --- | --- | --- | --- | --- | --- | --- |
| *E. coli* | MagnaPure Compact | 6 | 993 (2082) | 9.6 | *E. coli* (350) | 4.6 |
| *E. coli* | Qiagen Mini-column | 2 | 25847 (28191) | 326.3 | *E. coli* (24654) | 11.6 |
| *K. pneumoniae* | MagnaPure Compact | 48 | 914 (6602) | 15.3 | *K. pn* (117) | 2.3 |
| *K. pneumoniae* | Qiagen Mini-column | 24 | 43953 (48021) | 541.7 | *K. pn* (39082) | 11.3 |

Detection of antimicrobial resistance genes was carried out on selected data by using the Antimicrobial resistance application (AMRA). The *K. pneumoniae* mini-column sequence data described in Table 1 was subjected to further investigation by uploading on AMRA via the cloud-based Epi2Me agent. The strain is known to contain resistance genes, including the beta lactamase OKP-B-7 gene and the plasmid pKQPS2 resistance genes aadB (aminoglycoside), sul1 (sulfonamide), and ESBL encoding genes OXA-2 and SHV-18 (14). The report contained limited information, with 120 resistance gene hits, with various confidences and bacterial species. For *K. pnemoniae* AMRA only reported OKP-B-7 and SHV-18. These results apparently reflect the relatively high error rate (10-20%) in MinION sequencing reads. The report is available when logging in to Methrichor.

Next, we decided to attempt co-detection of 5 bacterial pathogens, within one sample, at relatively low concentrations ranging from as little as 5 ng DNA and downwards in a 10-log series, and also to test a re-used MinION flow cell. Because the MinION SQK-RAD003 protocol requires an input of at least 400 ng DNA, corresponding to a quantity as high as approximately 10^8^ bacterial cells, and because we are seeking a direct-from-clinical-sample protocol with no DNA amplification step, we furthermore decided to add each low concentration sample to 400 ng DNA from the low-pathogen *A. baumanii* to function as carrier. The DNA sequencing of this metagenomic sample did not give satisfactory results. During a 7 hr run, only 6280 reads were generated with a total yield of 102.6 Mb. None of the low concentration microbes were identified by WIMP, only *A. baumanii* (5070 reads). A sudden drop in available pores in the flow cell was observed as the experiment started, from 1003 down to 251 available pores. A separate experiment with a mixture of *A. baumanii* and *P. aeruginosa* provided similar disappointing results. Because our previous experiment related in Table 1 was performed with a naïve flow cell, the results were suggestive to us that flow cell reuse is associated with a significant and unacceptable drop in sensitivity and data accumulation even with DNA isolated on mini-columns.

Next, using manually mini-column extracted DNA from 4 different bacteria and using a naïve flow cell, we wanted to analyze a truly metagenomic sample. A high quantity (292 ng) of *P. mirabilis* DNA served as carrier, and was mixed with lower amounts of *H. influenzae* (105 ng), *K. pneumoniae* (52 ng) and *E. coli* (10 ng corresponding to approximately 2×10^6^ bacterial cells). A SQK-RAD003 sequencing library was constructed from the pool of DNA showing high purity (OD 260/280 ~1.8 and OD 260/230 2.0-2.2).

All four species were accurately identified (Figure 1), and the read numbers corresponded well with the genome content for each bacterium in the sequencing library; indicating that MinION can both qualitatively and quantitatively provide correct metagenomic data. Furthermore, there was a trend towards a relatively high read number as compared with colony count at the lower quantity range, suggesting a MinION sensitivity approaching clinically relevant concentrations at the ng level (corresponding to approximately 10^5^ bacterial cells); in agreement with the data reported recently on orthopedic device infection samples (9). These results indicate that a gentle manual DNA extraction procedure and single-use flow cells are critical parameters.

**Figure 1.**
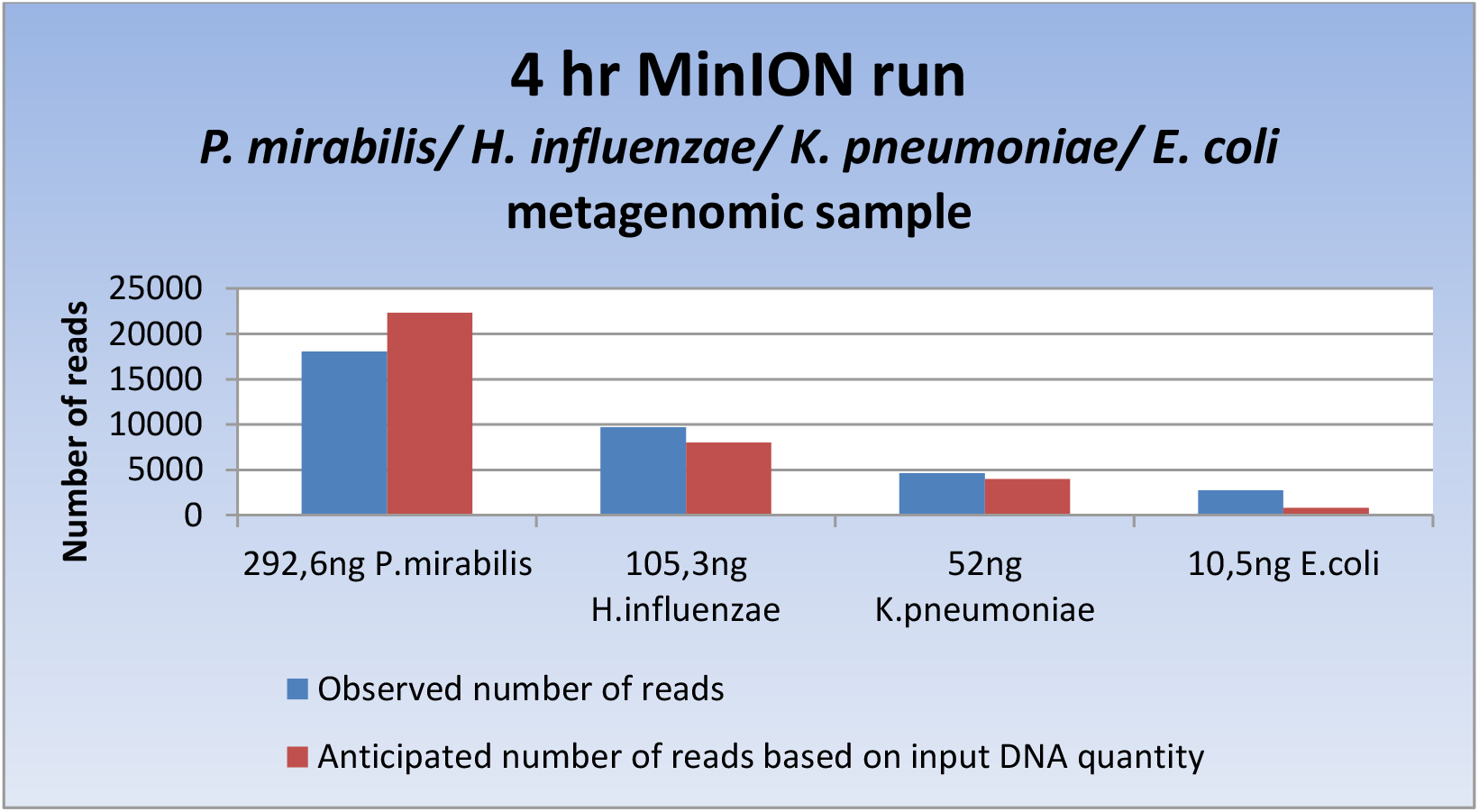
DNA sequencing results from a metagenomic sample containing *P. mirabilis, H. influenzae*, *K. pneumoniae* and *E. coli*.

Our combined preliminary assessment of MinION suggests to us that it readily identifies multiple microorganisms in a metagenomic manner. First and foremost MinION promises to become able to function as a rapid and accurate pathogen species identification technique for clinical specimens. The technology is under continued development, and refinement of sample preparation will most probably pose a major challenge, with a need to customize it according to sample type.

